# Using a handful of transcriptomes to detect sex-linked markers in a lizard with homomorphic sex chromosomes

**DOI:** 10.1101/2023.03.14.532509

**Authors:** Paul A. Saunders, Carles Ferre-Ortega, Peta Hill, Oleg Simakov, Tariq Ezaz, Christopher P. Burridge, Erik Wapstra

**Affiliations:** Discipline of Biological Sciences, University of Tasmania, Private Bag 5, Sandy Bay, Tasmania 7000, Australia; Institute for Applied Ecology, University of Canberra, Bruce, Australian Capital Territory 2601, Australia; Department of Molecular Evolution and Development, University of Vienna, Vienna 1010, Austria

**Keywords:** Reptile, *Carinascincus*, Sex determination, method, RNA-seq, PCR

## Abstract

To understand the biology of a species it is often crucial to be able to differentiate males and females. Many species lack distinguishable sexually dimorphic traits, but in those that possess sex chromosomes, molecular sexing offers a good alternative. Designing molecular sexing assays is typically achieved through the comparison of male and female genomic sequences, often from reduced-representation sequencing. However, in many non-model species sex chromosomes are poorly differentiated, and identifying sex-limited sequences and developing PCR-based sexing assays is challenging without additional genomic resources. Here we highlight a simple procedure for detection of sex-linked markers based on transcriptomes that circumvents limitations of other approaches. We apply it to the spotted snow skink *Carinascincus ocellatus*, a lizard with homomorphic XY chromosomes that also experiences environmentally-induced sex reversal. With transcriptomes from 3 males and 3 females alone, we identify thousands of putative Y-linked sequences. We confirm linkage through alignment of assembled transcripts to a distantly related genome, and readily design PCR primers to sex *C. ocellatus* and related species. In addition to providing an important molecular sexing tool for these species, this approach also facilitated valuable comparisons of sex determining systems on a large taxonomic scale.

## Introduction

Separate sexes are found in most species of animals and in many plants (including ~20% of crop species)^1,2^. Because males and females typically differ in traits such as morphology, behavior, and dispersal^3^, accurate sex identification is a crucial component of ecological, evolutionary, and conservation biology studies^4–7^. Unfortunately, identification of sex is challenging in numerous species lacking readily distinguishable sexually dimorphic phenotypes. For species in which sex is determined genetically, molecular sexing offers a solution, and has facilitated key outcomes for applied and fundamental research. For instance, the development of sex markers for birds^8^ induced a surge in empirical studies focused on sex allocation^9^.

The most rapid, reliable and affordable way to genetically sex individuals is through polymerase chain reaction (PCR). The development of sex-specific PCR primers requires the identification of sex-linked sequences, which can be challenging in species with homomorphic (*i.e.*, poorly differentiated) sex chromosomes^10^, which includes many animals outside of mammals and birds^11^. Currently, sex-linked sequences are typically identified through the analysis of next-generation DNA sequencing data, using a range of sampling strategies and bioinformatics workflows^12–16^. The most common approach for species with homomorphic sex chromosomes relies on reduced-representation sequencing (RRS) of multiple phenotypically sexed individuals, and the identification of sex-limited loci or variants. This approach is more affordable than whole genome sequencing, and has proved successful across a wide-range of organisms^14,17–30^. However, it has some drawbacks. Small sample sizes (<5-10 males and females) generate many false positives that are hard to identify without additional genomic resources^13,17,19,22^, and the typically short DNA reads from RRS also provide limited space for the design of diagnostic PCR primers^17,31^.

In this study we use a straightforward and cost-effective approach based on RNA-seq data that circumvents the requirements of large sample sizes to accurately detect sex-linked markers, and more readily generates PCR markers than other approaches. We applied this to identify sex-linked markers in the spotted snow skink, *Carinascincus ocellatus.* Recent research in this species using RRS (DArT-seq) identified homomorphic sex chromosomes (XY), and environmentally-induced sex reversal (XX males) in controlled conditions^32^. Given naturally occurring sex reversal is a rare feature in lizards^33^, a more thorough characterization of the sex-linked region(s) and development of a PCR sexing assay has crucial implications for understanding the evolution of sex determination in this species and potentially other squamates. Here, Based on RNA-seq data from 3 males and 3 females only, we identified hundreds of candidate male-limited sequences. Without additional genomic resources for *C. ocellatus*, we inferred sex-linkage of these sequences via the alignment of transcripts to the reference genome of a distantly related specie*s*. Assembled transcript sequences enabled easy development of several PCR primer sets, allowing high-throughput genotypic sexing in the spotted snow skink and related species. In addition to outlining a simple and efficient approach to identify and study homomorphic sex chromosomes, we introduce valuable genetic resources (transcriptomes and molecular sexing tools) for an emerging model system with respect to sex determination, sex allocation and sex reversal^32,34–37^.

## Methods

### Animals and RNA sequencing

Three males and females of *C. ocellatus* (Scinidae) were captured on Tasmania’s Central Plateau (41 51’S, 146 34’E) and sacrificed following methods approved by the University of Tasmania Animal Ethics Committee (Approval A23626). Brains and gonads were immediately dissected and snap frozen in liquid nitrogen. mRNA was extracted from the 12 samples at the Garvan Institute, and strand-specific RNA-seq libraries were generated and sequenced at the Ramaciotti Centre for Genomics, producing 127–185 million 100 nt paired-end reads per sample. Read quality was checked with FastQC v0.11.8, and quality trimming and adaptor clipping performed using Trimgalore v0.6.4 (default parameters; Table S1).

### Transcriptome assembly

Female samples were pooled to build a *de novo* transcriptome filtering out contigs shorter than 300 bp (using *Trinity v2.8.4* with options *–SS_lib_type RF–min_kmer_cov 2 – min_contig_length 300).* The resulting assembly was filtered using the Trinity scripts *align_and_estimate_abundance.pl* (with *Kallisto* alignment) and *filter_low_expr_transcripts.pl* to keep only the most highly expressed isoform for each gene (Table S2). A BUSCO analysis (v3.0.2; vertebrata database) revealed a considerable drop in redundancy after filtering, from 73% to 0.02% of duplicated genes among all complete genes identified (5.3% missing and 4.8% fragmented genes post-filtering).

### Detection and distribution of sex-limited variants

Trimmed reads of both sexes were mapped to the filtered female transcriptome using BWA-mem (0.7.16a-r1181). Reads with a mapping quality <20 were removed (samtools; v1.15), and duplicates marked using Picard *MarkDuplicates* (v2.21.4). Given that *C. ocellatus* has a male heterogametic (XY) sex determining system^38,39^, our aim was to identify sites where females are consistently homozygous and males consistently heterozygous. Variants (SNPs and short indels) were called for each sample using GATK’s *HaplotypeCaller* and resulting files were joined by sex using *GenomicsDBImport* and *GenotypeGVCFs.* GATK’s *VariantFiltration* was used on male variants to mark genotypes with a depth of coverage <3 as uncalled (-G-filter “DP<3” –set-filtered-genotype-to-no-call) and keep sites heterozygous in all or all but one sample (-filter “AN<8” and -filter “AF<0.35 || AF>0.65”), and on female variants to keep sites homozygous in all or all but one sample (-G-filter “DP<3” –set-filtered-genotype-to-no-call and -filter “AF>0.08). GATK’s *SelectVariants* was run to remove sites at which more than one sample was uncalled (--max-nocall-fraction 0.2), and bcftools (v1.12) was used to keep only biallelic sites. Finally, male and female variant files were merged using Bedtools (v2.29.2) *intersect* with *-u* option to keep filtered sites called in both sexes, homozygous in females and heterozygous in males, and therefore carrying male-limited variants.

If male-limited variants are Y-linked, the transcripts carrying them must originate from the non-recombining region of the X chromosome (as the reference transcriptome was built based on female (XX) samples). Transcripts carrying male-limited variants were aligned to the whole genome of the common wall lizard *Podarcis muralis* using *Blastn* (v. 2.12.0), and clustering of best hits within the reference was used to identify potentially Y-linked genes in *C. ocellatus.* This approach assumes some level of synteny, but no requirement of homology of sex determining regions among species.

### Approach validation

To provide an indication of false positive error rate, we ran the whole workflow on the same RNA-seq reads but assuming a ZW sex determining system (reference transcriptome assembled with male reads, search for female-limited SNPs: homozygous in males and heterozygous in females). To verify the method is applicable to other species with homomorphic sex chromosomes, we also ran the workflow on a publicly available dataset: brain and gonad RNA-seq from 3 males and 3 females of the African cichlid *Gnathochromis pfefferi*, which was recently shown to have poorly differentiated sex chromosomes, homologous to *Oreochromis niloticus* linkage groups 11 and 15^40^ (accession numbers: SRR9688100, SRR9688103, SRR9688131, SRR9688134, SRR9688136, SRR9688139, SRR9688465, SRR9688466, SRR9688473, SRR9688477, SRR9688578, SRR9688583). Given this cichlid dataset is around 10 times smaller than ours (average of 1.2 Gbp of data per sample), this also provided the opportunity to assess the suitability of the approach to datasets from lower sequencing effort.

### Designing and testing PCR primers for sex-linkage

Transcripts with male-limited variants that aligned in clusters within the *P. muralis* genome were manually scanned to find regions suitable for the design of sex diagnostic PCR assays in *C. ocellatus.* We specifically targeted regions on transcripts carrying several variants within approximately 20 bp (size of a PCR primer). Triplet PCR primer sets were designed using Primer3 and Oligoanalyzer (https://eu.idtdna.com/calc/analyzer). Each triplet is composed of two “outer” primers (one forward and one reverse) that should hybridize to both the X and Y, and one “inner” primer (forward or reverse), that overlaps the male specific variants and should hybridize only to the Y (Table S3). This way, electrophoresis of a single PCR can discriminate XX and XY individuals (one *vs.* two bands).

The primer triplets were tested on previously sampled *C. ocellatus* with known phenotypic and genotypic sex^32^: five males and females with concordant genotypic and phenotypic sex, and five ‘sex reversed’ XX males. All individuals were born in captivity from females captured pregnant in the wild on the Tasmanian Central Plateau (41 59’S, 146 44’E). Tests were also run against 15 individuals from a genetically distant and previously sampled coastal population (42°34’ S, 147°52’ E)^32,41^. Cycling conditions were 95 °C for 3 min followed by 34 cycles of 95 °C for 30s (denaturation), 56 °C for 30s (annealing; may vary based on primer sets, Table S3), 72 °C for 1 min (extension), and then a final 72 °C extension for 5 min. PCR products were visualized using 1.5% agarose gel electrophoresis dyed with Midori Green Advance. PCR products were Sanger sequenced to confirm identity to the target region.

Finally, the primer sets were tested on samples of known sex from four other *Carinascincus (C. pretiosus, C. metallicus, C. greeni* and *C. microlepidotus*) and two other more distantly related skinks (*Pseudomoia entrecasteauxii* and *Liopholis whitii*). Known male and female *C. ocellatus* were included as positive controls when testing the other species.

## Results

### Identification of male-limited variants

The comparison of male and female RNA-seq datasets identified 5,082 male-limited variants distributed across 831 transcripts. Homologous sequences for 70% (582/831) of these transcripts were identified in the common wall lizard *Podarcis muralis*, and a clear enrichment (74.5% of all aligned transcripts) was observed along chromosome 10 (Fig. 1A). Most variants clustered in three distinct regions (Fig. 2, Table S4). The first and largest region (2,276 variants) is in the middle of Pmu10 (28–41 Mb). The two other regions are more distal (63–64.3 Mb and 70.2–75.3 Mb) and contain less variants (230 and 503, respectively). In parallel, we identified 1,395 female-limited variants, distributed across 728 transcripts. While homologous sequences were found in *P. muralis* for a similar proportion of these transcripts containing female-limited variants (73%, 533/728), they appeared randomly distributed in the reference (Fig. 1B, Table S5).

**Figure 1.**
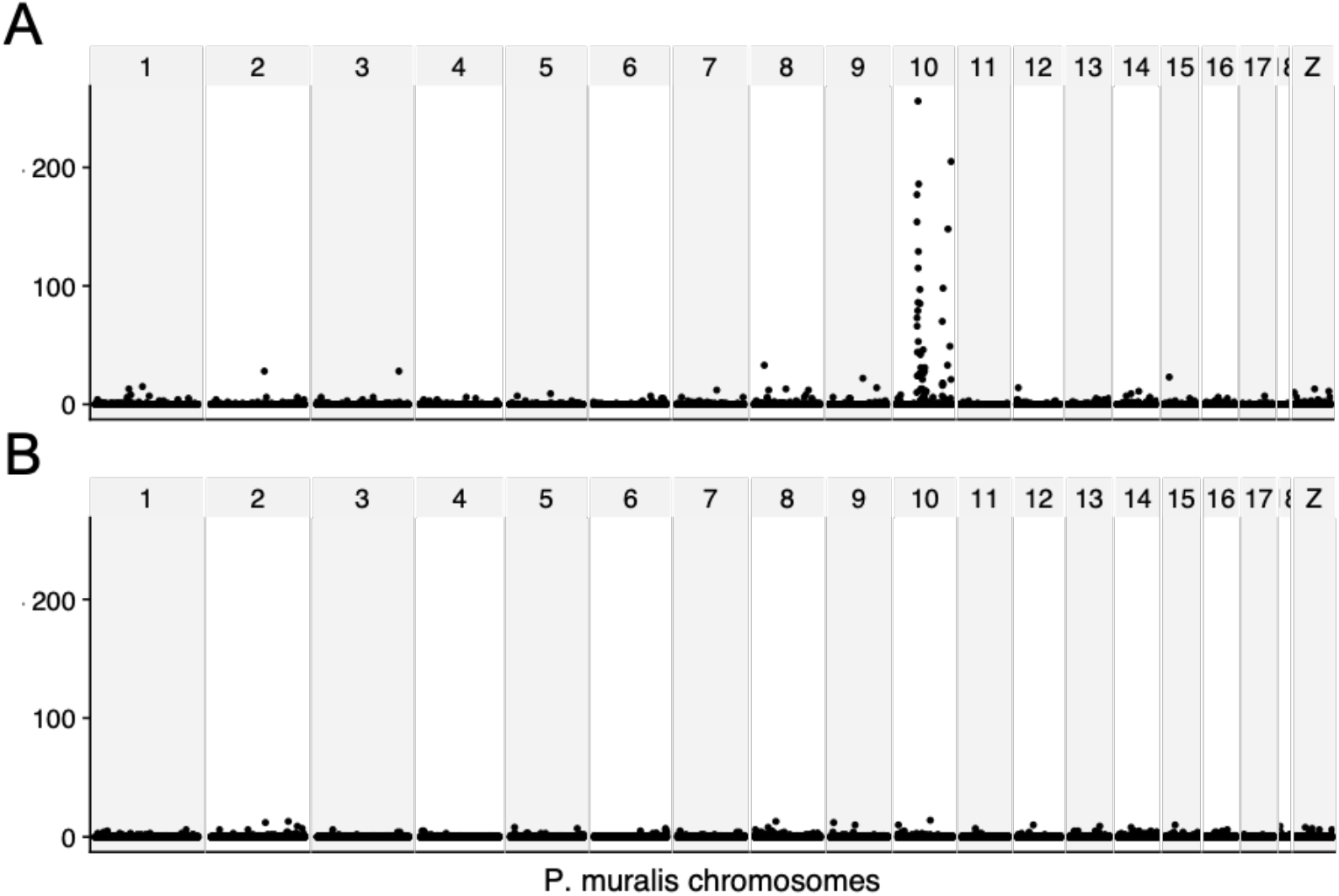
Distribution of sex-limited variants in *C. ocellatus* transcripts based on their mapping position along the chromosomes of *P. muralis.* A. male-limited variants. B. female-limited variants. Each dot represents the number of variants in a window of 100 kbp.

**Figure 2.**
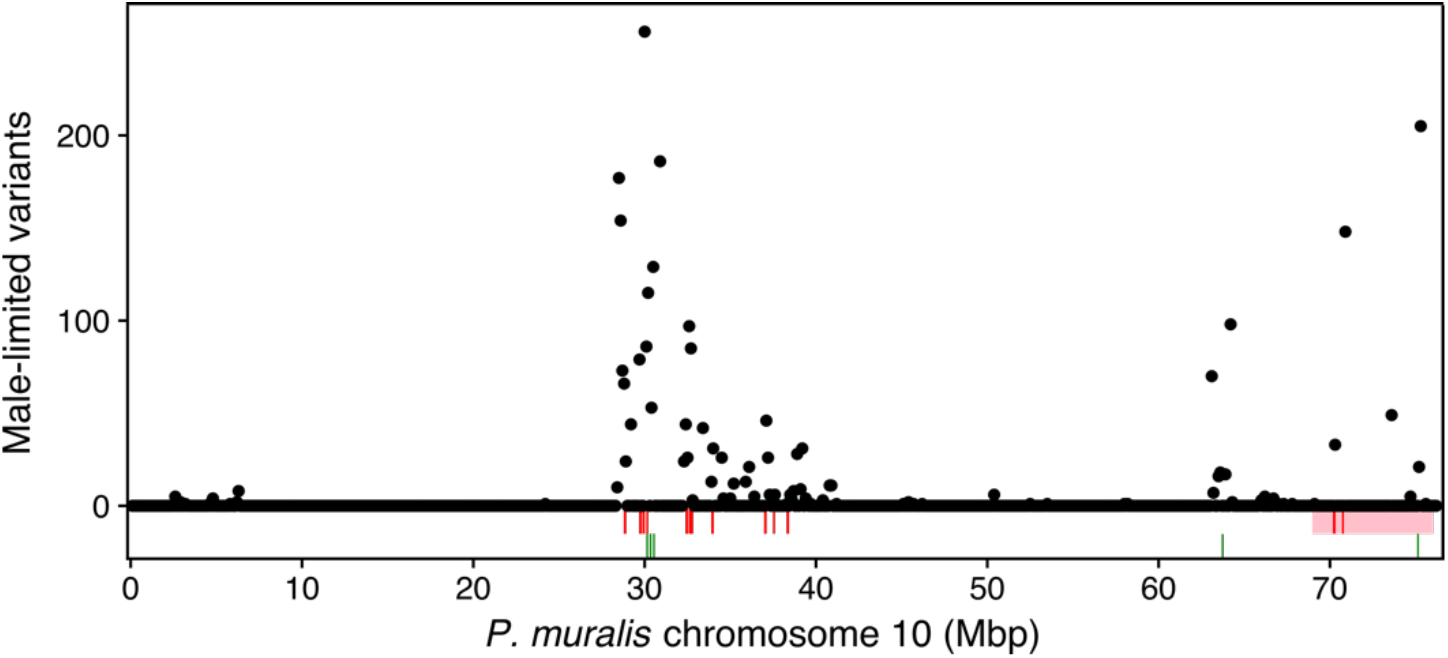
Distribution of male-limited variants in *C. ocellatus* transcripts based on their mapping position along *Podarcis muralis* chromosome 10 (Pmu10). Each dot represents the number of variants in a window of 100 kbp. The pink area shows the region assumed to be sex-linked across multiple other skink species, identified by Kostmann et al.^47^. Red bars show the position of orthologs of sex-linked genes in *Eulamprus heatwolei*^44^. Green bars show the position of transcripts used to design the PCR primer sets (see table S3).

We tested the validity of our approach by analyzing RNA-seq data from the African cichlid *Gnathochromis pfefferi.* We identified 1,463 male-limited variants scattered across 449 transcripts. >99% of transcripts mapped to the *O. niloticus* reference genome, and >80% of variants aligned to LG11 and LG15 (Fig. S1) as expected.

### PCR assays

Five *C. ocellatus* transcripts were selected to develop PCR primers to (i) independently validate their sex linkage and (ii) provide efficient molecular sexing markers (Table S3). All five aligned to Pmu10, in the three distinct regions showing enrichment in male-limited variants (Fig. 2). In *C. ocellatus*, all primer sets produced two bands in known XY males, and one band in known XX females and sex-reversed XX males, in both the Central Plateau and the coastal populations (Table 1, fig. S2). This confirms sex linkage of the markers and reproducibility of the PCR molecular sexing. Sanger sequencing of randomly selected PCR products in multiple males and females confirmed amplification of targeted fragments. Except for primer set 5, DNA was always successfully amplified in the other species tested, reflecting at least partial sequence conservation (Table 1, fig. S3). The first three primer sets distinguished sexes across the other *Carinascincus* tested, except for primer set 1 in *C. greeni* and *C. microlepidotus.* In the two more distantly-related species *L. whitii* and *P. entrecasteauxii*, no primer set worked as intended.

**Table 1.**
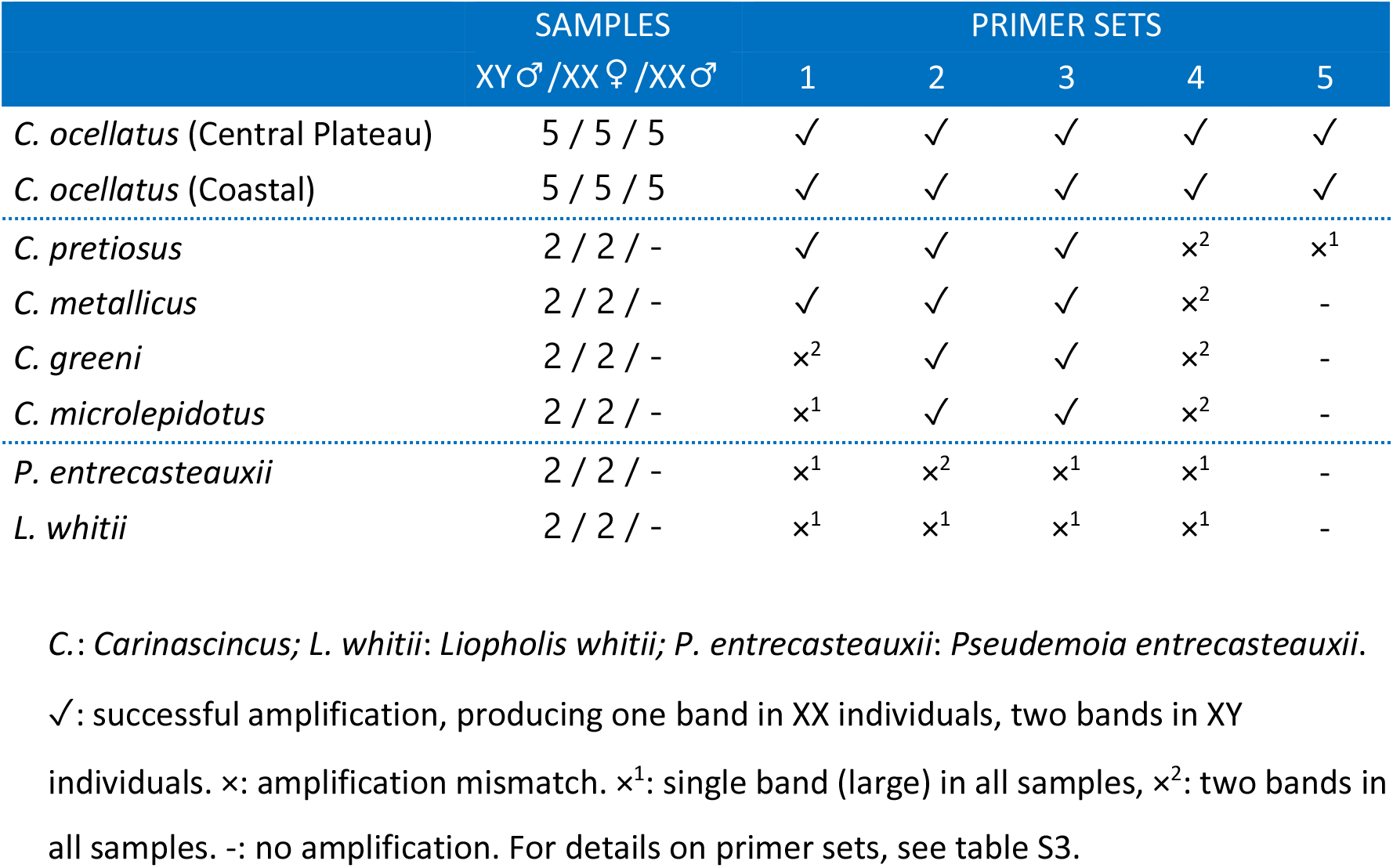
PCR tests across multiple *Carinascincus* and other skinks species.

## Discussion

Using RNA-seq data for only 3 males and 3 females and a reference genome from a distantly related species (different infraorder), we identified several thousand putative male-limited variants in the spotted snow skink *C. ocellatus.* From these we easily developed molecular sexing PCR tests, validating Y-linkage and allowing a simple and high throughput genotypic sexing of *C. ocellatus* and related species. We validated our bioinformatic approach by re-analysing a smaller dataset from the cichlid fish *Gnathochromis pfefferi*, which was also recently shown to have poorly differentiated sex chromosomes^40^ (Fig. S1). The use of RNAseq circumvents some limitations of reduced-representation sequencing (RSS) for detection of sex-linked sequences in non-model species, yet still shares the positives of being affordable, fast and easy to implement, and allowing the identification of sex-linked markers even in species with very young sex chromosomes (a few million years or less^19^).

The key advantages of using RNAseq are facilitated by the longer lengths of assembled transcripts (often thousands of base pairs) relative to the short unassembled reads of RRS (usually 50–150 bp). This greatly facilitates assessments of sequence homology to distantly related reference genomes (i.e., when the target species lacks an assembled genome). This in turn enables the identification of sex-linked regions based on the clustering of candidate sex-linked markers derived from only a small number of individuals. In contrast, the alignment of RRS reads (usually done with BLAST) tends to have very poor success when no close reference genome is available (1–10%^20,25,26,42,43^), such that there is a stronger reliance on large sample sizes to increase confidence on whether candidate markers are sex-linked. This was the case for sex-linkage inference of *C. ocellatus* DArT-seq markers, as only 2% of the approximately 300 sex-linked markers aligned to BLAST’s nucleotide collection^38,44^. In contrast, over 70% of our assembled transcripts carrying putative male-limited variants successfully aligned to the wall lizard genome despite an estimated divergence time of 174Mya^45^. This greater mapping rate is decisive when only a handful of individuals are sequenced as it circumvents the high false-positive error rates of sex-linkage when based on genotype frequencies alone. This was also demonstrated by our testing for female-limited variants, consistent with a ZW system. Although we identified ~1400 putative female-limited variants based on the analysis of 3 males and 3 females, their transcripts were did not cluster within the reference genome, arguing against sex-linkage.

Ultimately, true sex-linkage should be confirmed via large scale sexing analyses, and another advantage of the longer assemblies derived from transcriptome is the easy development of PCR assays, even when no other genomic resources are available. Reads in most RRS studies are simply too short to design PCR sexing assays^18,38^. Loci detected as sex-limited based on RRS restriction-site presence-absence also often fail PCR validation; they tend to amplify in both sexes when restriction-sites are flanked by sequences homologous on the X and Y (or Z and W)^31,46^. With RNA-seq data, primers can be designed based on the assembled transcripts, providing much more information to develop reliable PCR assays. In our experiment, all 5 transcripts selected to design PCR assays could successfully discriminate male and female spotted snow skinks.

Based on the alignment of the aforementioned transcripts to the wall lizard genome, we show that that the sex-determining region in *C. ocellatus* is homologous to median and terminal regions of *P. muralis* chromosome 10 (Fig. 2). Recent studies in the water skink *Eulamprus heatwolei* ^44^ and the common sandfish *Scincus scincus* ^47^ also identified sex-linked markers with homology to Pmu10. Sex-linked markers in *E. heatwolei* align mostly to the median region of Pmu10, overlapping with the position where most sex-linked variants in *C. ocellatus* align (Fig. 2). In contrast, the markers identified in *S. scincus* align to the terminal region of Pmu10 (69–76Mbp), with minimal overlap with a few sparsely distributed candidate sex-linked *C. ocellatus* variants. This discrepancy could reflect differences in genomic regions which are sex-linked among species, or different sensitivities in the approaches used in each study. Indeed, while we used a ‘SNP-density’ approach, best suited to detect regions of sex chromosomes with little differentiation, the two other studies rely on ‘subtraction’ approaches, effective only if there is substantial sequence divergence between the sex chromosomes^48^. It has been suggested that the exact XY sex determination system identified in *S. scincus* is widely conserved across all major skink lineages based on outcomes from qPCR sexing in 12 other species^49^. Hypothetically, *C. ocellatus* might share with other skinks an ‘old and diverged’ sex determining region homologous to 69–76 Mbp of Pmu10, while also harbouring a less divergent, more recently sex-linked region homologous to Pmu10 28–40 Mbp.

The amplification of several of our sexing markers across four other *Carinascincus* species suggest that the sex determining system of *C. ocellatus* is shared across the genus. However, the fact that all markers failed in two more distantly related skinks casts doubt on the conservation of sex determination across all Scincidae, as previously suggested based on work in the Egernia subfamiliy and in the ground skink *Scincella melanostica^25,50^.* Testing the PCR markers developed here and by others^25,44,51^ across a larger panel species would benefit our understanding of scincid lizard sex chromosome evolution. More broadly, our study highlights the benefits of increasing the number of available sex-linked genetic markers and molecular sexing tests in taxonomic groups with poorly studied sex determination.

## Supplementary material

**Figure S1**. Distribution of male-limited variants in *Gnathochromis pfefferi* transcripts based on their mapping position along *Oreochromis niloticus.*

**Figure S2**. Validation of sex-linkage using PCR across multiple males and females of *C. ocellatus.*

**Figure S3**. Molecular sexing of male and female samples in other species.

**Table S1.** Raw reads quality check and trimming statistics.

**Table S2.** Transcriptome assemblies statistics.

**Table S3.** Details on PCR primer sets.

**Table S4.** Male-limited transcripts and alignment to *Podarcis muralis* genome.

**Table S5.** Female-limited transcripts and alignment to *Podarcis muralis* genome.

## References

1. Leonard, J. L. The Evolution of Sexual Systems in Animals. Transitions Between Sexual Systems (2018). doi:10.1007/978-3-319-94139-4_1.

2. Renner, S. S. The relative and absolute frequencies of angiosperm sexual systems: Dioecy, monoecy, gynodioecy, and an updated online database. Am. J. Bot. 101, 1588–1596 (2014).

3. Trivers, R. Parental investment and sexual selection. in *Sexual Selection and the Descent of Man* 136–79 (B Campbell, 1972).

4. Benestan, L. et al. Sex matters in massive parallel sequencing: Evidence for biases in genetic parameter estimation and investigation of sex determination systems. Mol. Ecol. 26, 6767–6783 (2017).

5. Ancona, S., Dénes, F. V., Krüger, O., Székely, T. & Beissinger, S. R. Estimating adult sex ratios in nature. Philos. Trans. R. Soc. B Biol. Sci. 372, (2017).

6. Mossman, C. A. & Waser, P. M. Genetic detection of sex-biased dispersal. Mol. Ecol. 8, 1063–1067 (1999).

7. Robertson, B. C., Elliott, G. P., Eason, D. K., Clout, M. N. & Gemmell, N. J. Sex allocation theory aids species conservation. Biol. Lett. 2, 229–231 (2006).

8. Fridolfsson, A. A. & Ellegren, H. A Simple and Universal Method for Molecular Sexing of Non-Ratite Birds. J. Avian Biol. 30, 116–121 (1999).

9. Komdeur, J. & Pen, I. Adaptive sex allocation in birds: The complexities of linking theory and practice. Philos. Trans. R. Soc. B Biol. Sci. 357, 373–380 (2002).

10. Palmer, D. H., Rogers, T. F., Dean, R. & Wright, A. E. How to identify sex chromosomes and their turnover. Mol. Ecol. 28, 4709–4724 (2019).

11. Bachtrog, D. et al. Sex Determination: Why So Many Ways of Doing It? PLoS Biol. 12, e1001899 (2014).

12. Muyle, A. et al. Sex-detector: A probabilistic approach to study sex chromosomes in non-model organisms. Genome Biol. Evol. 8, 2530–2543 (2016).

13. Käfer, J., Lartillot, N., Marais, G. A. B. & Picard, F. Detecting sex-linked genes using genotyped individuals sampled in natural populations. Genetics 218, (2021).

14. Feron, R. et al. RADSex: A computational workflow to study sex determination using restriction site--associated DNA sequencing data. Mol. Ecol. Resour. 21, 1715–1731 (2021).

15. Nursyifa, C., Brüniche-Olsen, A., Garcia-Erill, G., Heller, R. & Albrechtsen, A. Joint identification of sex and sex-linked scaffolds in non-model organisms using low depth sequencing data. Mol. Ecol. Resour. 22, 458–467 (2022).

16. Sigeman, H., Sinclair, B. & Hansson, B. Findzx: an automated pipeline for detecting and visualising sex chromosomes using whole-genome sequencing data. BMC Genomics 23, 1–14 (2022).

17. Gamble, T. et al. Restriction Site-Associated DNA Sequencing (RAD-seq) Reveals an Extraordinary Number of Transitions among Gecko Sex-Determining Systems. Mol. Biol. Evol. 32, 1296–309 (2015).

18. Liu, H. et al. Sex-specific markers developed by next-generation sequencing confirmed an XX/XY sex determination system in bighead carp (Hypophthalmichthys nobilis) and silver carp (Hypophthalmichthys molitrix). DNA Res. 25, 257–264 (2018).

19. Jeffries, D. L. et al. A rapid rate of sex-chromosome turnover and non-random transitions in true frogs. Nat. Commun. 9, 1–11 (2018).

20. Brelsford, A., Lavanchy, G., Sermier, R., Rausch, A. & Perrin, N. Identifying homomorphic sex chromosomes from wild-caught adults with limited genomic resources. Mol. Ecol. 17, 752–759 (2017).

21. Mathers, T. C. et al. Transition in sexual system and sex chromosome evolution in the tadpole shrimp Triops cancriformis. Heredity 115, 37–46 (2015).

22. Scharmann, M., Ulmar Grafe, T., Metali, F. & Widmer, A. Sex is determined by xy chromosomes across the radiation of dioecious nepenthes pitcher plants. Evol. Lett. 3, 586–597 (2019).

23. Gamble, T. & Zarkower, D. Identification of sex-specific molecular markers using restriction site-associated DNA sequencing. Mol. Ecol. Resour. 14, 902–913 (2014).

24. Gamble, T. et al. The Discovery of XY Sex Chromosomes in a Boa and Python. Curr. Biol. 27, 2148–2153 (2017).

25. Bouffet-Halle, A. et al. Characterisation and cross-amplification of sex-specific genetic markers in Australasian Egerniinae lizards and their implications for understanding the evolution of sex determination and social complexity. Aust. J. Zool. 69, 33–40 (2022).

26. Nielsen, S. V et al. Escaping the evolutionary trap ? Sex chromosome turnover in basilisks and related lizards (Corytophanidae: Squamata). Biol. Lett. 15, 20190498 (2019).

27. Nielsen, S. V, Banks, J. L., Diaz Jr., R. E., Trainor, P. A. & Gamble, T. Dynamic sex chromosomes in Old World chameleons (Squamata: Chamaeleonidae). J. Evol. Biol. 31, 484–490 (2018).

28. Nielsen, S. V., Daza, J. D., Pinto, B. J. & Gamble, T. ZZ/ZW sex chromosomes in the endemic puerto rican leaf-toed gecko (*Phyllodactylus wirshingi*). Cytogenet. Genome Res. 157, 89–97 (2019).

29. Keating, S. E. et al. Sex chromosome turnover in bent-toed geckos (Cyrtodactylus). Genes 12, 1–11 (2021).

30. Keating, S. E., Griffing, A. H., Nielsen, S. V., Scantlebury, D. P. & Gamble, T. Conserved ZZ/ZW sex chromosomes in Caribbean croaking geckos (Aristelliger: Sphaerodactylidae). J. Evol. Biol. 33, 1316–1326 (2020).

31. Fowler, B. L. S. & Buonaccorsi, V. P. Genomic characterization of sex-identification markers in Sebastes carnatus and Sebastes chrysomelas rockfishes. Mol. Ecol. 25, 2165–2175 (2016).

32. Hill, P. et al. Sex reversal explains some, but not all, climate mediated differences in sex ratio within a viviparous reptile. Proc. R. Soc. B 286, 20220689 (2022).

33. Whiteley, S. L., Castelli, M. A., Dissanayake, D. S. B., Holleley, C. E. & Georges, A. Temperature-Induced Sex Reversal in Reptiles: Prevalence, Discovery, and Evolutionary Implications. Sex. Dev. 15, 148–156 (2021).

34. Pen, I. et al. Climate-driven population divergence in sex-determining systems. Nature 468, 436–438 (2010).

35. Gruber, J., Cunningham, G. D., While, G. M. & Wapstra, E. Disentangling sex allocation in a viviparous reptile with temperature-dependent sex determination: a multifactorial approach. J. Evol. Biol. 31, 267–276 (2018).

36. Cunningham, G. D., While, G. M. & Wapstra, E. Climate and sex ratio variation in a viviparous lizard. Biol. Lett. 13, 0–3 (2017).

37. Wapstra, E., Uller, T., While, G. M., Olsson, M. & Shine, R. Giving offspring a head start in life: Field and experimental evidence for selection on maternal basking behaviour in lizards. J. Evol. Biol. 23, 651–657 (2010).

38. Hill, P. L., Burridge, C. P., Ezaz, T. & Wapstra, E. Conservation of sex-linked markers among conspecific populations of a viviparous skink, *Niveoscincus ocellatus*, exhibiting genetic and temperature-dependent sex determination. Genome Biol. Evol. 10, 1079–1087 (2018).

39. Hill, P., Shams, F., Burridge, C. P., Wapstra, E. & Ezaz, T. Differences in Homomorphic Sex Chromosomes Are Associated with Population Divergence in Sex Determination in *Carinascincus ocellatus*. Cells 10, 291 (2021).

40. El Taher, A., Ronco, F., Matschiner, M., Salzburger, W. & Böhne, A. Dynamics of sex chromosome evolution in a rapid radiation of cichlid fishes. Sci. Adv. 7, (2021).

41. Cliff, H. B., Wapstra, E. & Burridge, C. P. Persistence and dispersal in a Southern Hemisphere glaciated landscape: The phylogeography of the spotted snow skink (*Niveoscincus ocellatus*) in Tasmania. BMC Evol. Biol. 15, 1–13 (2015).

42. Nielsen, S. V., Pinto, B. J., Guzmán-Méndez, I. A. & Gamble, T. First report of sex chromosomes in night lizards (Scincoidea: Xantusiidae). J. Hered. 111, 307–313 (2020).

43. Keating, S. E., Greenbaum, E., Johnson, J. D. & Gamble, T. Identification of a cis-sex chromosome transition in banded geckos (Coleonyx, Eublepharidae, Gekkota). J. Evol. Biol. 35, 1675–82 (2022).

44. Cornejo-Páramo, P. et al. Viviparous reptile regarded to have temperature-dependent sex determination has old XY chromosomes. Genome Biol. Evol. 12, 924–930 (2020).

45. Kumar, S., Stecher, G., Suleski, M. & Blair Hedges, S. TimeTree: A Resource for Timelines, Timetrees, and Divergence Times. Mol. Biol. Evol. 34, 1812–1819 (2017).

46. Gamble, T. Using RAD-seq to recognize sex-specific markers and sex chromosome systems. Mol. Ecol. 25, 2114–2116 (2016).

47. Kostmann, A., Kratochvíl, L. & Rovatsos, M. Poorly differentiated XX/XY sex chromosomes are widely shared across skink radiation. Proc. R. Soc. B Biol. Sci. 288, 20202139 (2021).

48. Cortez, D. et al. Origins and functional evolution of Y chromosomes across mammals. Nature 508, 488–93 (2014).

49. Thépot, D. Sex chromosomes and master sex-determining genes in turtles and other reptiles. Genes 12, (2021).

50. Patawang, I. et al. Cytogenetics of the skinks (Reptilia, Scincidae) from Thailand; IV: newly investigated karyotypic features of *Lygosoma quadrupes* and *Scincella melanosticta*. Caryologia 71, 29–34 (2018).

51. Dissanayake, D. S. B. et al. Identification of Y chromosome markers in the eastern three-lined skink (*Bassiana duperreyi*) using in silico whole genome subtraction. BMC Genomics 21, (2020).

