## Supplementary Figures S1-S3 for "Using a handful of transcriptomes to detect sex-linked markers in a lizard with homomorphic sex chromosomes"

**Figure S1**. Distribution of male-limited variants in *Gnathochromis pfefferi* transcripts based on their mapping position along *Oreochromis niloticus.*

**Figure S2**. Validation of sex-linkage using PCR across multiple males and females of *C. ocellatus.*

**Figure S3**. Molecular sexing of male and female samples in other species.

**Figure S1**

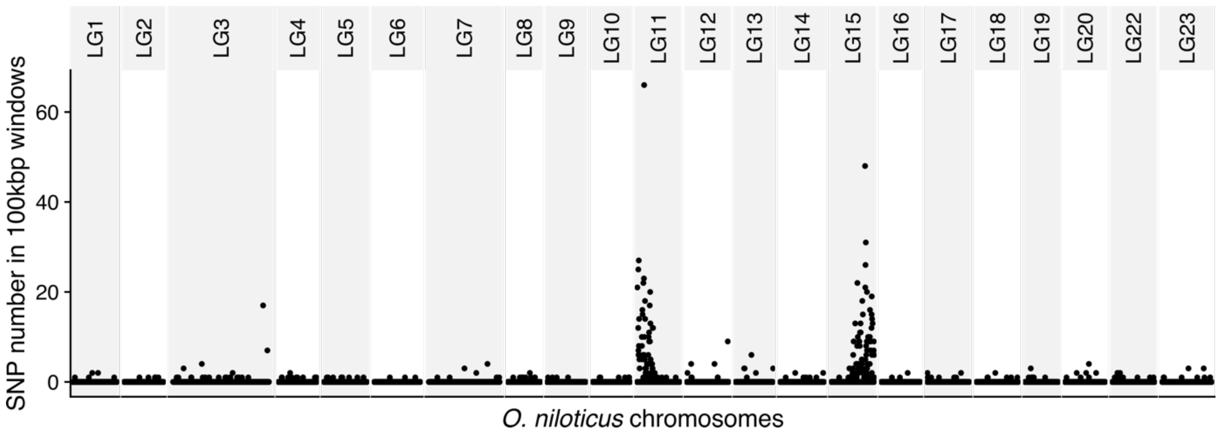
LG1
LG2
LG3
LG4
LG5
LG6
LG7
LG8
LG9
LG10
LG11
LG12
LG13
LG14
LG15
LG16
LG17
LG18
LG19
LG20
LG22
LG23
SNP number in 100kbp windows
60
40
20
0
*O. niloticus* chromosomes

**Figure S2**

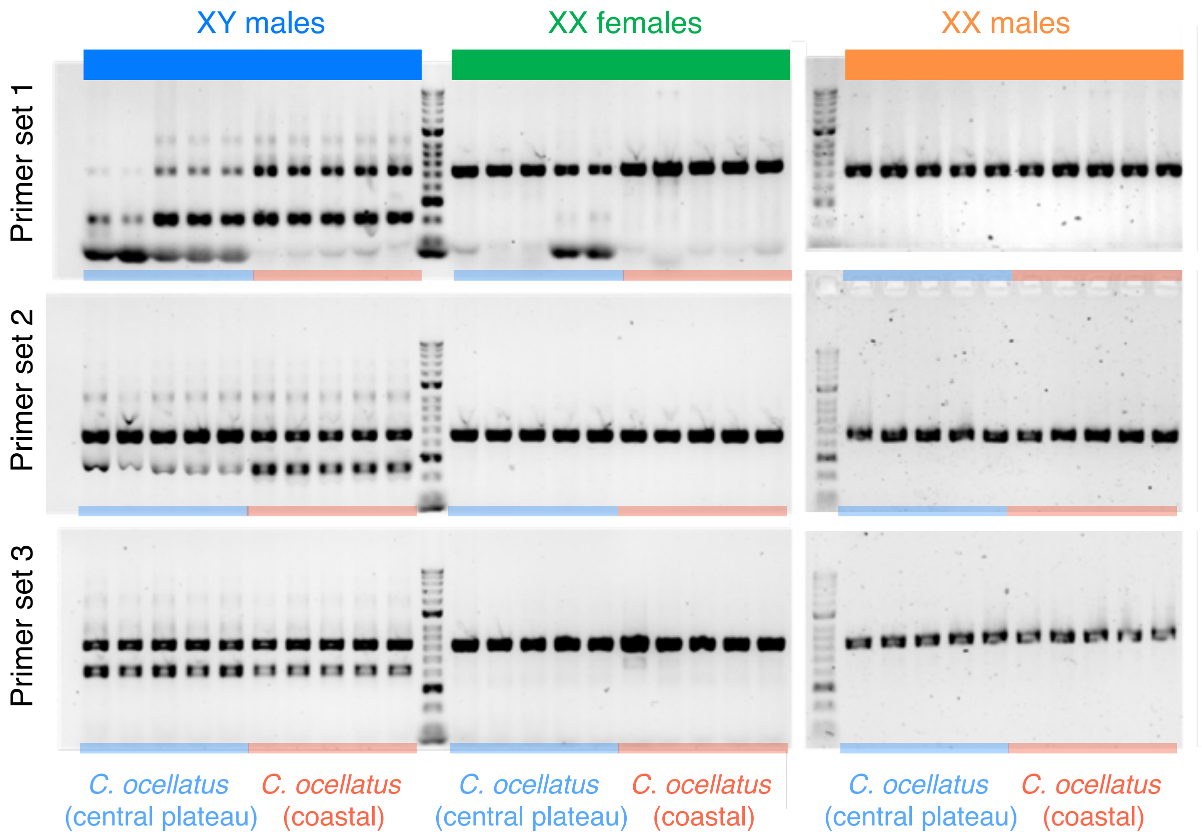

XY males
XX females
XX males
Primer set 1
Primer set 2
Primer set 3
C. ocellatus (central plateau)
C. ocellatus (coastal)
C. ocellatus (central plateau)
C. ocellatus (coastal)
C. ocellatus (central plateau)
C. ocellatus (coastal)

### Figure S3

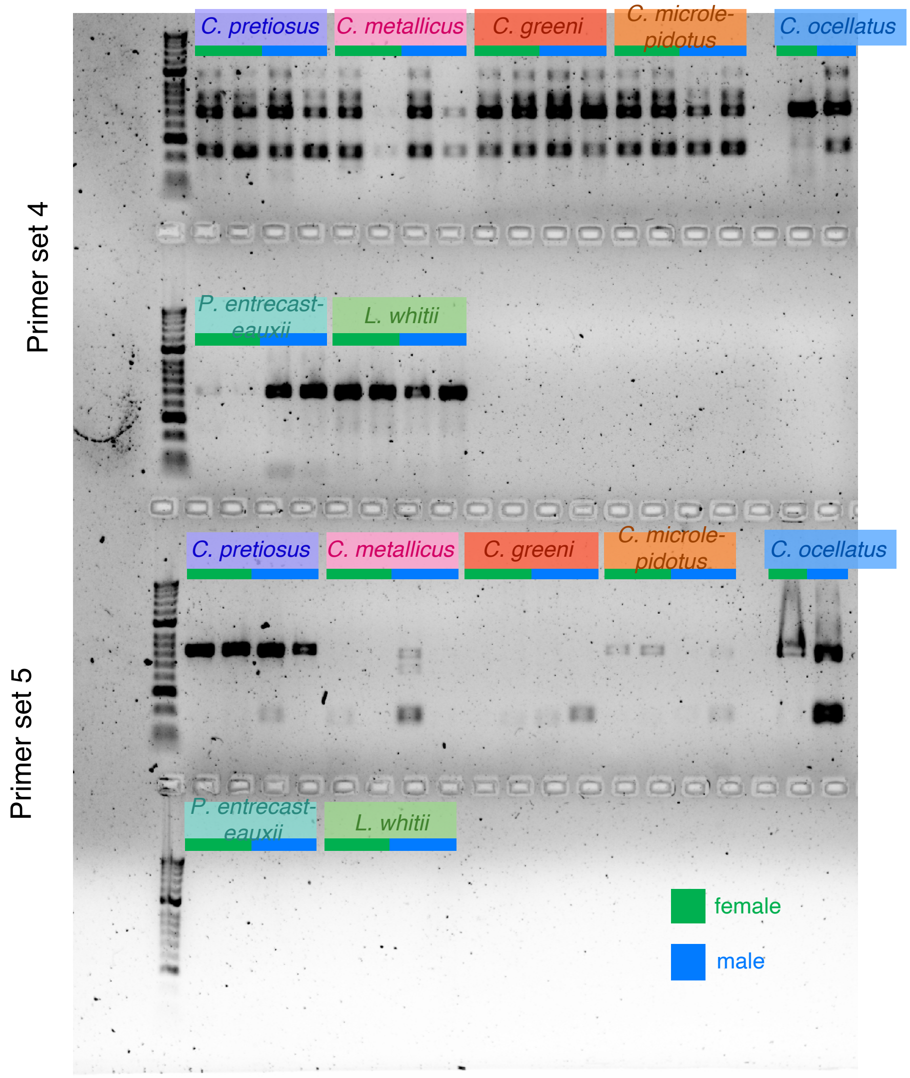
Primer set 4
Primer set 5
C. pretiosus
C. metallicus
C. greeni
C. microle-
pidotus
C. ocellatus
P. entrecast-
eauxii
L. whitii
female
male
